# Vitamin D receptor protects against dysbiosis and tumorigenesis via the JAK/STAT pathway in intestine

**DOI:** 10.1101/2020.02.18.946335

**Authors:** Yong-Guo Zhang, Rong Lu, Shaoping Wu, Ishita Chatterjee, David Zhou, Yinglin Xia, Jun Sun

## Abstract

**Background:** Vitamin D exerts regulatory roles via vitamin D receptor (VDR) in mucosal immunity, host defense, and inflammation involving host factors and microbiome. Human *Vdr* gene variation shapes the microbiome and VDR deletion leads to dysbiosis. Low VDR expression and diminished vitamin D/VDR signaling are observed in colon cancer. Nevertheless, how intestinal epithelial VDR is involved in tumorigenesis through gut microbiota remains unknown. We hypothesized that intestinal VDR protects mice against dysbiosis via modulating the JAK/STAT pathway in tumorigenesis. To test our hypothesis, we used an azoxymethane/Dextran Sulfate Sodium-induced cancer model in intestinal VDR conditional knockout (VDR^ΔIEC^) mice, cell cultures, stem-cell derived colonoids, and human colon cancer samples.

**Results:** VDR^ΔIEC^ mice have higher numbers of tumors with location shifted from distal to proximal colon. Fecal microbiota analysis showed that VDR deletion leads to bacterial profile shift from normal to susceptible carcinogenesis. We found enhanced bacterial staining in mouse and human tumors. Microbial metabolites from VDR^ΔIEC^ mice showed elevated secondary bile acids, consistent with the observations in human CRC. We further identified that VDR protein bound to the Jak2 promoter, suggesting that VDR transcriptionally regulated Jak2. The JAK/STAT pathway is critical in intestinal and microbial homeostasis. Fecal samples from VDR^ΔIEC^ mice activate the STAT3 activation in human and mouse organoids. Lack of VDR led to hyperfunction of Jak2 in respond to intestinal dysbiosis. A JAK/STAT inhibitor abolished the microbiome-induced activation of STAT3.

**Conclusion:** We provide insights into the mechanism of VDR dysfunction leading to dysbiosis and tumorigenesis. It indicates a new target — microbiome and VDR for prevention of cancer.

## Background

Current research has implicated vitamin D deficiency as a critical factor in the pathology and clinical outcome of colon rectal cancer (CRC) [1, 2]. Low plasma vitamin D is associated with adverse CRC survival after surgical resection [3, 4]. Vitamin D receptor (VDR) is a nuclear receptor that mediates functions of 1,25-dihydroxyvitamin D (1,25(OH)_2_D_3_), the biological active form of vitamin D [5]. Higher VDR expression in tumor stromal fibroblast is associated with longer survival in a large cohort of CRC patients [2]. The parallel appreciation of a role for the VDR in cancer biology began approximately 3 decades ago and subsequently a remarkable increase has occurred in the understanding of its actions in normal and malignant systems [6].

The VDR regulation of gut microbiome in human and animal studies represents a newly identified and highly significant activity for VDR [7–9]. Human *Vdr* gene variation shapes gut microbiome and *Vdr* deletion leads to dysbiosis [8]. Our study on VDR and bacteria establishes a microorganism-induced program of epithelial cell homeostasis and repair in the intestine [10]. Dysregulation of bacterial-host interaction can result in chronic inflammatory and over-exuberant repair responses, and is associated with the development of various human diseases including cancers [11, 12]. Even though vitamin D/VDR is an active topic in cancer research, the mechanism underlying host-microbiome interactions in cancer is incompletely understood. We know little about the mechanisms for the intestinal epithelial VDR and microbiome in CRC.

In the current study, we focused on the functions of VDR in intestinal epithelial cells and the microbiome. We hypothesized that intestinal VDR protects mice against dysbiosis via modulating the JAK/STAT pathway in tumorigenesis. VDR is required for intestinal epithelium functions and microbial homeostasis. We tested our hypothesis in an azoxymethane/Dextran Sulfate Sodium (AOM/DSS)-induced cancer model, using intestinal VDR conditional knockout VDR^ΔIEC^ mice, colonoids, and human samples. Lack of the VDR signaling pathway led to increased tumors in colon and shift tumor distribution in the intestinal VDR knockout (KO) mice. We investigated how the absence of intestinal VDR leads to dysfunction in epithelial cells-microbiome interactions and the mechanism through the JAK/STAT3 signaling. Emerging data suggest that interference JAK/STAT3 pathway may suppress the growth of colon cancer [13, 14]. JAK/STAT inhibitors are clinically used in patients with inflammatory bowel diseases [15]. Thus, VDR regulation of JAK/STAT3 pathway indicates a new target—microbiome and VDR signaling in anti-inflammation and anti-cancer. Our study provides new insights into the mechanisms of VDR in maintaining intestinal and microbial homeostasis and protecting against intestinal tumorigenesis.

## Results

### Intestinal epithelial VDR KO mice have higher tumor numbers and shifted tumor location

We tested our hypothesis in an AOM/DSS-induced cancer model using intestinal epithelial VDR conditional knockout VDR^ΔIEC^ mice (Fig. 1a). AOM mice develop hyperproliferative colonic mucosa, aberrant crypt foci (ACF), and eventually carcinomas [16]. AOM-DSS provides a widely used paradigm to study colitis-associated colon cancer. There was a striking difference in tumor incidence in mice with VDR^LoxP^ and VDR^ΔIEC^ mice. We found the VDR^ΔIEC^ mice developed more tumors (Fig. 1b and c). The number and size of tumors were significantly bigger in the VDR^ΔIEC^ mice compared with the VDR^LoxP^ mice (Fig. 1c and d). Interestingly, tumor location in the VDR^ΔIEC^ mice significantly shifted from distal to proximal colon, compared to tumors mainly in the distal colon of VDR^LoxP^ mice (Fig. 1b and e). Furthermore, the pathological analysis of colon samples (Fig. 1f) indicated difference of tumor stage (carcinoma versus adenoma) between VDR^ΔIEC^ mice and VDR^LoxP^ AOM/DSS experimental groups. Epithelial hyperproliferation plays a critical role in the development of colon cancer. Our IHC data of proliferative marker PCNA showed that PCNA in colon was significantly increased in the VDR^ΔIEC^ mice, compared to the VDR^LoxP^ mice (Fig. 1g).

**Fig. 1.**
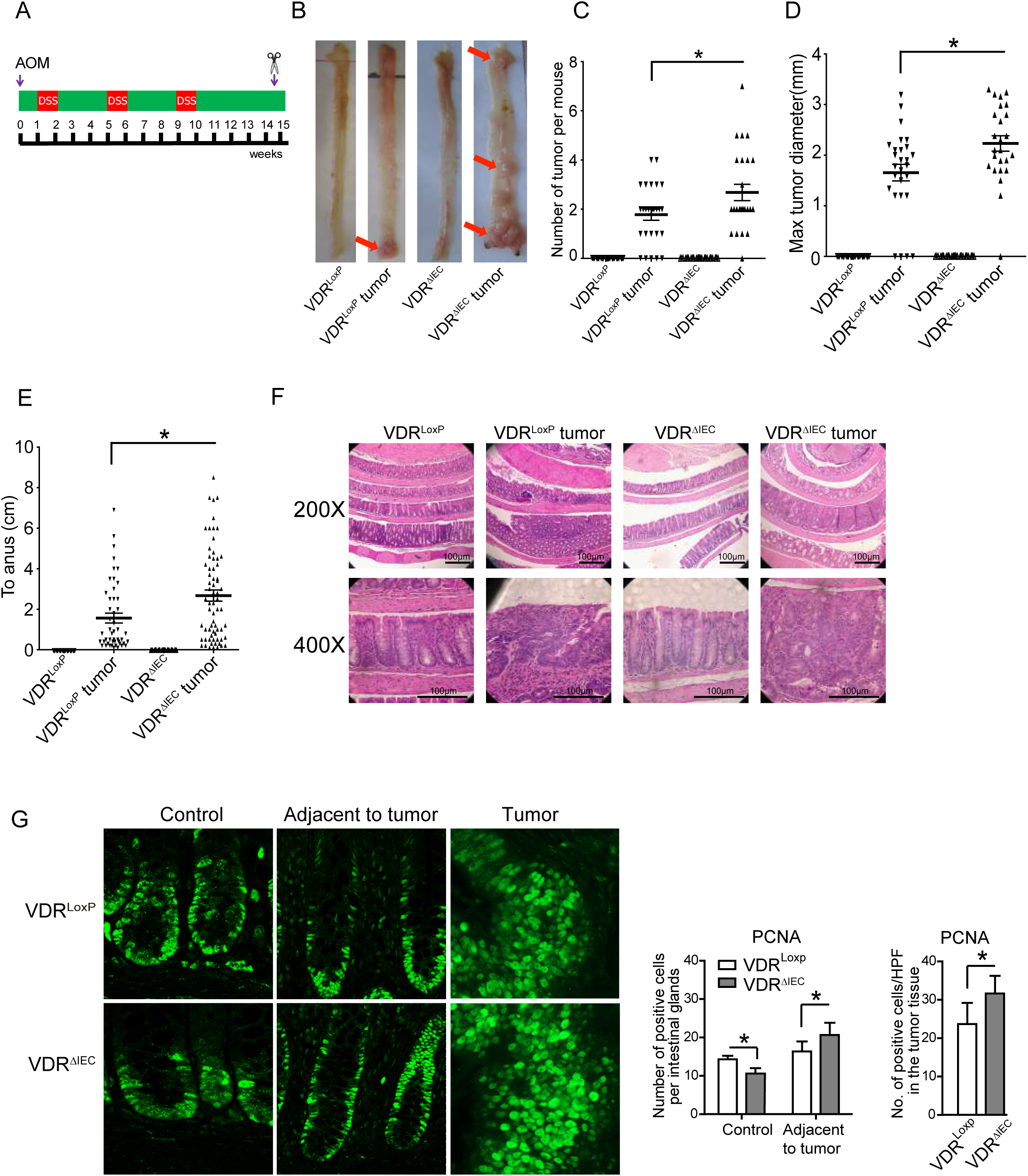
Intestinal epithelial cell VDR KO mice developed more tumors. **a** Schematic overview of the AOM-DSS induced colon cancer model. AOM (10 mg/kg) was injected on day 0. At Day 7, 2% DSS solution was administered to mice in drinking water. Seven days of DSS is followed by three weeks of drinking water. An additional two cycles of DSS were administered prior to sacrification. At Week 15, mice were sacrificed. **b** Colonic tumors in situ. Representative colons from different groups. Tumors were indicated by red arrows. **c** Tumor numbers in AOM-DSS induced colon cancer model: VDR^LoxP^ and VDR^ΔIEC^ mice. Data are expressed as mean ± SD. n = 25-30, one-way ANOVA test, *P < 0.05. No tumors in controls for VDR^LoxP^ and VDR^ΔIEC^ mice, therefore controls are not included for comparisons. **d** Max tumor size in AOM-DSS induced colon cancer model: VDR^LoxP^ and VDR^ΔIEC^ mice. (Data are expressed as mean ± SD. n = 25-30, one-way ANOVA test, *P < 0.05. **e** The distance of each tumor to the anus was measured. (Data are expressed as mean ± SD. n = 25-30, one-way ANOVA test, *P < 0.05. **f** Representative H&E staining of “Swiss rolls” of representative colons from the indicated groups. Images are from a single experiment and are representative of 10 mice per group. **g** Quantitation of PCNA-positive cells in control mucosa/per intestinal glands or in the tumors tissue/high-power field. PCNA expression in the tumor tissue of VDR^ΔIEC^ mice was significantly higher than that in the VDR^LoxP^ mice. Data are from a single experiment and are representative of 5 mice per group. (Data are expressed as mean ± SD. n = 5, student’s t-test, *P < 0.05).

### Lack of intestinal VDR leads to dysbiosis and shift of bacterial profile for the higher risk of CRC

VDR^ΔIEC^ mice lacking intestinal epithelial VDR is known to have dysbiosis [7]. Using 16S sequencing methods, we showed the difference of fecal microbiome between VDR^ΔIEC^ mice and VDR^LoxP^ mice (n=10 each) at the genus level (n=10) (Fig. 2a). Fig. 2b showed the Unweighted UniFrac distances of stool samples from VDR^LoxP^ and VDR^ΔIEC^ mice on a principal coordinate analysis (PCoA) scale. We further showed the percentages of the affected genera between VDR^LoxP^ mice and VDR^ΔIEC^ mice (Fig. 2c). Functional alterations of the intestinal microbiome were detected by fecal microbiota KEGG analysis. Lacking VDR leads to bacterial profile shift from normal to carcinogenesis susceptibility (Fig. 2d), indicating that cancer risk was significantly higher in the VDR^ΔIEC^ mice.

**Fig. 2.**
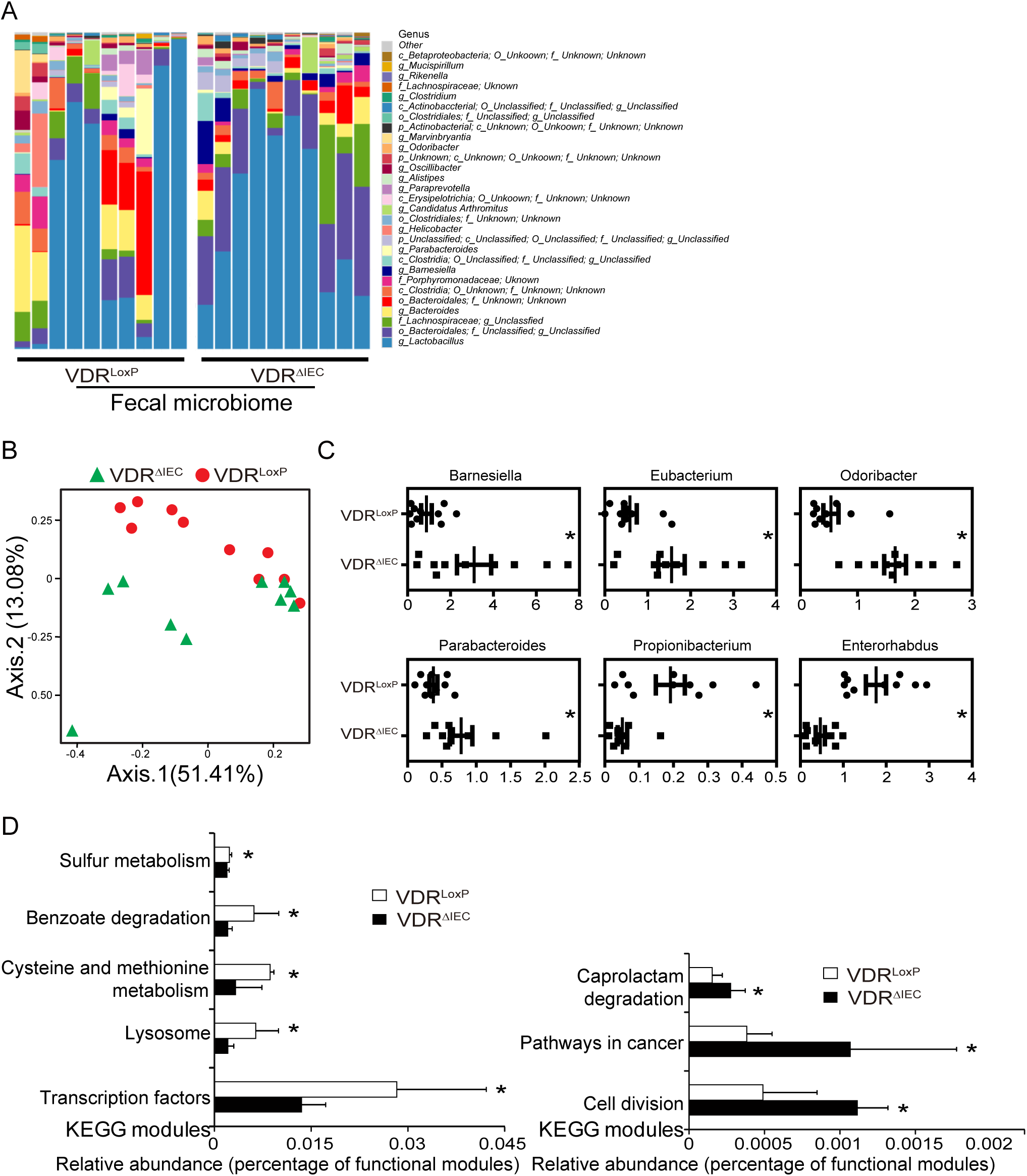
Dysbiosis leads to high cancer risk in the VDR^ΔIEC^ mice. **a** Composition of the bacterial community at the genus level in stool samples from separate cages of VDR^LoxP^ mice(n=10) and VDR^ΔIEC^ (n=10) mice. **b** Unweighted UniFrac distances of stool samples from VDR^LoxP^ and VDR^ΔIEC^ mice on a principal coordinate analysis (PCoA) scale. **c** The percentages of the affected genera were compared between VDR^LoxP^ mice and VDR^ΔIEC^ mice. (Data are expressed as mean ± SD. n = 10, Welch’s two-sample t-test, *P < 0.05). **d** Functional alterations of the intestinal microbiome related to vitamin D receptor (VDR) status. (Data are expressed as mean ± SD. n = 10, student’s t-test, *P < 0.05).

### VDR deletion enhanced bacteria in the tumors of VDRΔIEC mice and impacted bile acid metabolism

We then analyzed the relative bacteria abundance in the tumors. *Bacteroides fragilis,* a bacterial species enhanced in colon cancer, showed more staining in the tumors of VDR^ΔIEC^ mice, compared to the VDR^LoxP^ mice (Fig. 3a). Fig. 3b further showed that *Bacteroidales fragilis*, *Butyivibrio fibrisolvens* and *Firmicuyes peptostreptococcus* were enhanced in tumors in VDR^ΔIEC^ mice compared to VDR^LoxP^ mice in tumor tissue. These bacteria are known to be associated with changes of metabolite (e.g. short chain fatty acids, bile acids) in CRC [17–19].

**Fig. 3.**
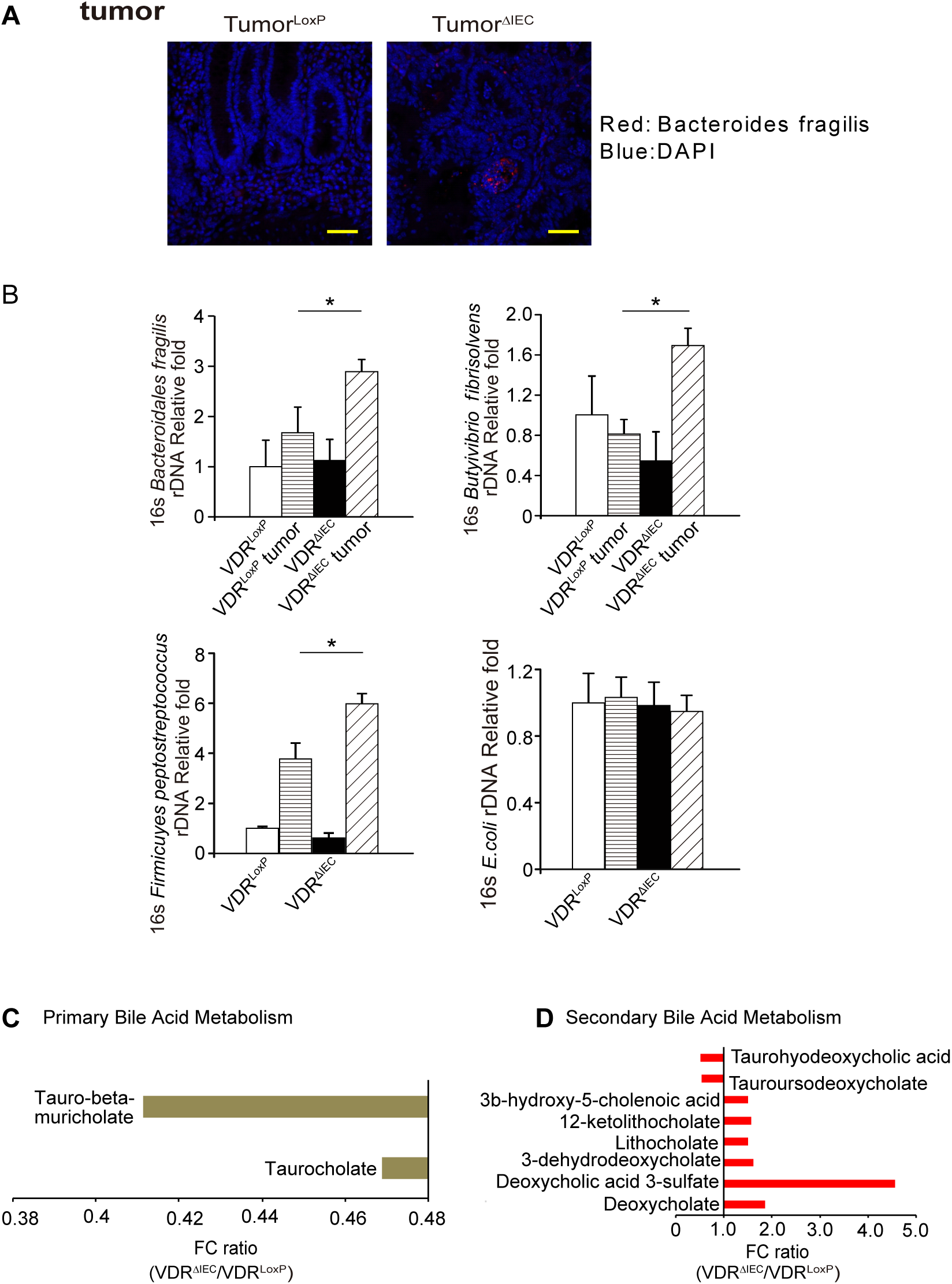
Lacking intestinal VDR leads to dysbiosis and shift of bacterial profile. **a** More *Bacteroides fragilis* in tumor tissue of VDR^ΔIEC^ mice were found by FISH. Images are from a single experiment and are representative of 5 mice per group. Scale bars, 40 µm. **b** *Bacteroidales fragilis*, *Butyivibrio fibrisolvens* and *Firmicuyes peptostreptococcus* were enhanced in tumors in VDR^ΔIEC^ mice compared to VDR^LoxP^ mice. (Data are expressed as mean ± SD. n = 6, one-way ANOVA test, *P < 0.05). **c** The fold change ratios of the average concentrations of primary bile acid in the VDR^ΔIEC^ group was significantly lower, compared to that in the control group. (VDR^LoxP^, n = 16; VDR^ΔIEC^, n = 17, Welch’s two-sample t-test, Metabolite ratio < 1.00, P < 0.05). **d** The fold change ratios of the average concentrations of secondary bile acid in the VDR^ΔIEC^ group was significantly higher, compared to that in the control group. (VDR^LoxP^, n = 16; VDR^ΔIEC^, n = 17, Welch’s two-sample t-test, Metabolite ratio ≥ 1.00, P < 0.05).

We quantitively profiled metabolites derived from host-microbial co-metabolism in fecal samples using the unbiased method. We found the changes in primary bile acid metabolism and secondary bile acid metabolism in VDR^ΔIEC^ mice. The fold change ratios of the identified bile acid species were significantly higher in the VDR^ΔIEC^ group than those in the control group (Fig. 3c and d).

These changes are consistent with the recent observations in human CRC that the bile acid metabolism is among the top biomarkers of patients [20].

### Increased inflammation in the VDR^ΔIEC^ mice

We further hypothesized that the altered intestinal epithelial and microbial functions lead to chronic inflammation, thus exacerbating colon cancer progression. We assessed several lymphocyte markers in normal colon and colonic tumors. Levels of CD68, CD3, and CD11b significantly increased in tumors, especially in VDR^ΔIEC^ mice (Fig. 4a). We also detected the cytokines in serum samples from VDR^LoxP^ and VDR^ΔIEC^ mice with or without tumor. We found that the level of FGF basic and MCP-1 in the tumor tissue of VDR^LoxP^ mice were higher than that of VDR^ΔIEC^ mice (Fig. 4b). In the gastrointestinal tract, tissue barrier integrity is particularly important. Serum samples from VDR^LoxP^ and VDR^ΔIEC^ mice were used to measure bacterial endotoxin with Limulus amebocyte lysate chromogenic endpoint assays. We found more bacterial endotoxin LPS in VDR^ΔIEC^ mice than in VDR^LoxP^ mice, especially in tumor groups (Fig. 4c). Lcn-2 is used as a marker of intestinal inflammation [21]. We found that the expression level of fecal Lcn-2 was significantly higher in tumor tissue of VDR^ΔIEC^ mice than that of the VDR^LoxP^ mice (Fig. 4d).

**Fig. 4.**
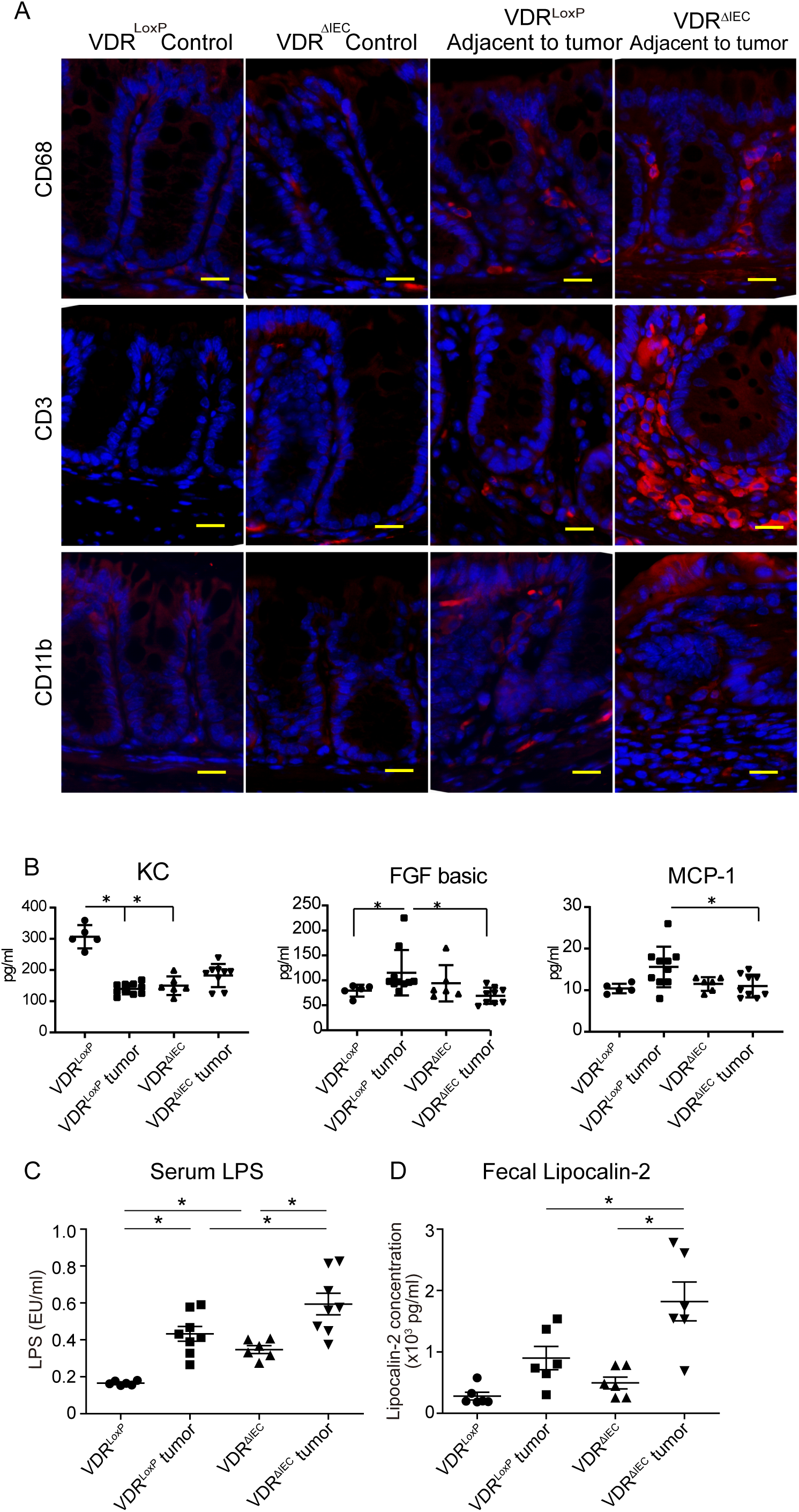
Altered intestinal epithelial and microbial functions may lead to chronic inflammation. **A** Several lymphocyte markers were detected in colon tissue by immunofluorescence staining. Levels of CD68, CD3, and CD11b significantly increased in tumors, especially in VDR^ΔIEC^ mice. Scale bars, 20 µm. **b** Serum samples were collected from VDR^LoxP^ and VDR^ΔIEC^ mice with or without tumor, then cytokines were detected by Luminex detection system. (Data are expressed as mean ± SD. n = 5-10, one-way ANOVA test, *P < 0.05). **c** Serum LPS was significantly high in the VDR^ΔIEC^ mice. (Data are expressed as mean ± SD. n = 6, one-way ANOVA test, *P < 0.05). **d** Fecal lipocalin-2 was increased in the VDR^ΔIEC^ mice with tumors. (Data are expressed as mean ± SD. n = 6, one-way ANOVA test, *P < 0.05)

### VDR deletion leads to hyperfunction of the Jak2 / STAT3 signaling in the tumor tissue

The JAK/STAT3 pathway is known to suppress the growth of colon cancer [13]. After the AOM/DSS treatment, in VDR^ΔIEC^ mice, we observed upregulated Jak2 and STAT3 proteins expression in colon cancer tissue using immunostaining (Fig. 5a). Further, Western blots confirmed Jak2 and STAT3 enhance expression tumors in AOM/DSS induced VDR^ΔIEC^ mice (Fig. 5b). But VDR deletion did change the expression of STAT1 and STAT5 in the colon tumor tissue (data not shown). Interestingly, without any treatment, VDR deletion led to reduced STAT3 and Jak2 in the basal level of cells at the protein level and mRNA level (Fig. 5c and 5d). Further, we identified that VDR protein bound to the Jak2 promoter (TGAACTTCTGAGAATTCA) by CHIP assay (Fig. 5e). Taken together, our observations show that absence of intestinal epithelial VDR leads to the hyperfunction of JAK/STAT3 signaling in inflammation.

**Fig. 5.**
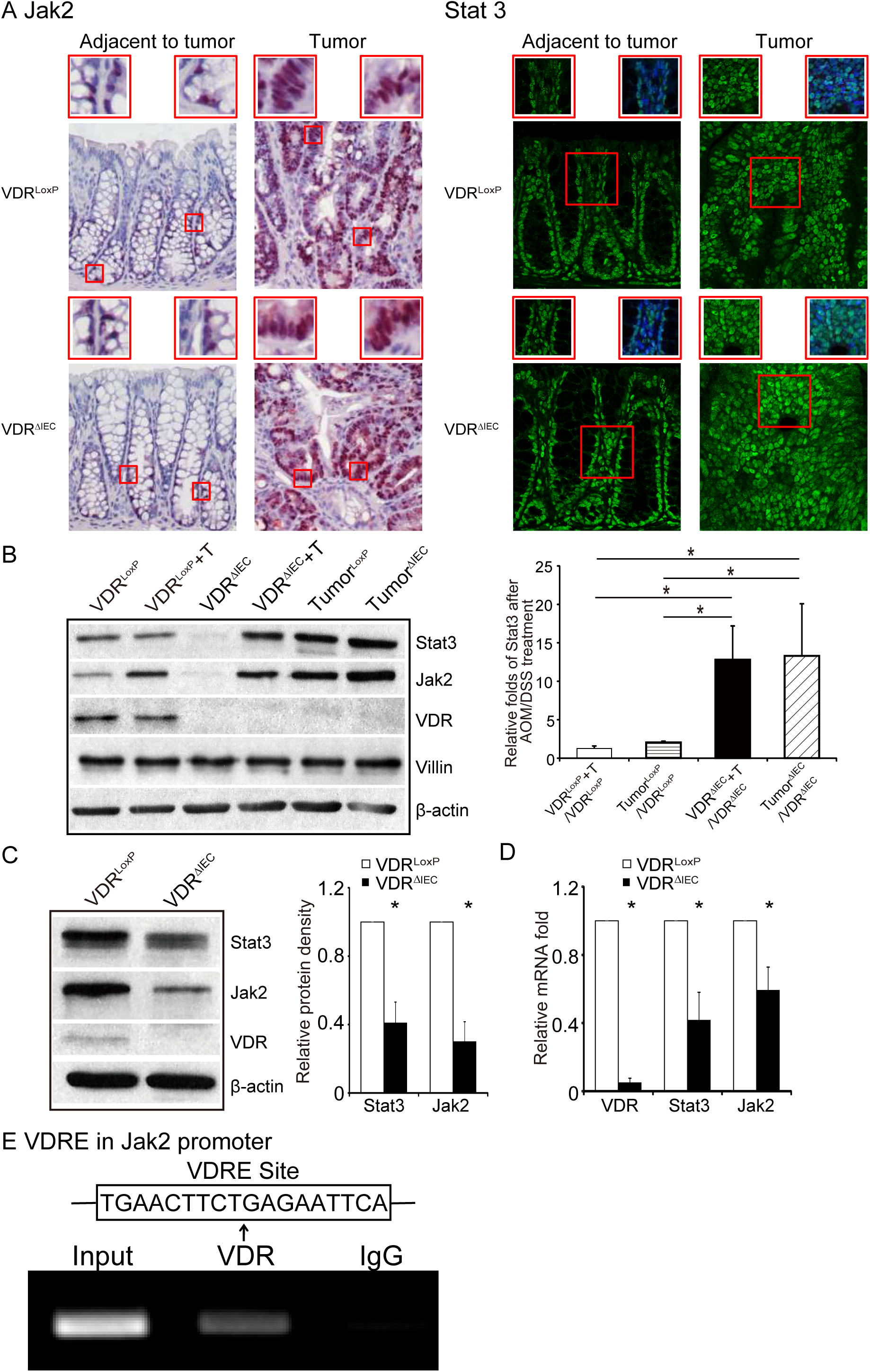
VDR deletion leads to dysfunction of the Jak2 / Stat3 signaling in the tumor tissue. **a** Jak2 and Stat3 were increased in tumor tissue of VDR^ΔIEC^ mice, compared to the tumor tissue of VDR^LoxP^ mice by immunofluorescence staining. Images are from a single experiment and are representative of 6 mice per group. **b** VDR deletion increased Jak2 and Sat3 in the colon tumor tissue. (Data are expressed as mean ± SD. n = 3, one-way ANOVA test, *P < 0.05). **c** VDR deletion decreased Jak2 and Stat3 at protein levels in colon. (Data are expressed as mean ± SD. n = 5, student’s t-test, *P < 0.05). **d** VDR deletion decreased Jak2 and Stat3 at mRNA levels in colon without any treatment. (Data are expressed as mean ± SD. n = 5, Welch’s two-sample t-test, *P < 0.05). **e** VDRE binds to the Jak2 promoter. CHIP-PCR amplification demonstrated binding of VDR to the promoter regions of Jak2. PCR were performed including input and negative controls. n = 3 separate experiments.

### Gut microbiome from VDRΔIEC mice actives the JAK/STAT signaling in colonoids

Using the stem-cell derived colonoids systems (Fig. 6a), we further investigated the influence of intestinal VDR during the activation of the JAK/STAT signaling. PCNA, a proliferation marker, and β-catenin were increased in the VDR^ΔIEC^ feces treated group followed by activation of stat3 (human colonoids in Fig. 6b). The similar hyperregulation of STAT3 was also observed in the mouse colonoids treated with microbiome from VDR^ΔIEC^ mice (Fig. 6c). We then treated the organoind with static, a STAT3 inhibitor. The total stat3 were decreased compare to the no-stattic treated mouse colonoids (Fig. 6d). However, the expressions of stat3 and β-catenin in the VDR^ΔIEC^ group were still higher than the VDR^LoxP^ group (Fig. 6d). We observed the similar effect of static in inhibiting the microbiome-activation of Jak2/ STAT3 signaling the mouse colonoids (Fig. 6e). Interestingly, stattic treatement also reduced the proliferation regulator β-catenin and the proliferation marker PCNA in colonoids.

**Fig. 6.**
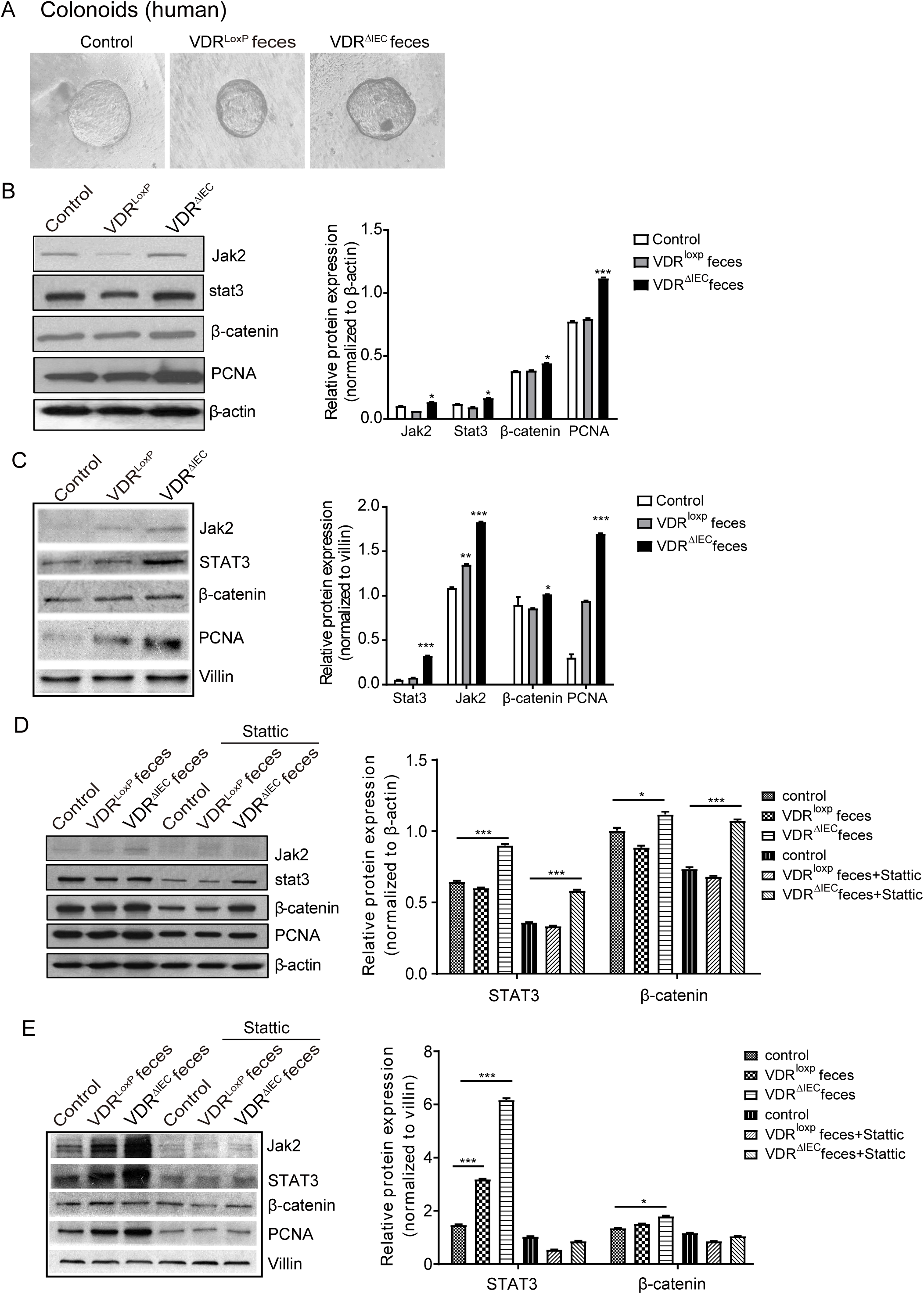
Gut microbiota from VDR^ΔIEC^ mice actives the JAK/STAT signaling in human and mouse organoids. **a** Human colonoids were prepared and treated with feces from VDR^lox^ or VDR^ΔIEC^ mice for 2 hours. **b** The expressions of Jak2 and Stat3 in human colonoids and **c** mouse colonoids were detected by western blots. PCNA and beta-catenin were increased in the VDR^ΔIEC^ feces treated group. Data are expressed as mean ± SD. n = 3, one-way ANOVA test, *P < 0.05, **P < 0.01, ***P < 0.001 compare to the control group. **d** Human and **e** mouse organoids were pretreated with 20 µM of stattic for 2h, then treated with feces for 2 hours. The expression of Jak2 and was increased after static treated, especially in the VDR^ΔIEC^ group. The total stat3 were decreased compare to the no-stattic treated group. Data are expressed as mean ± SD. n = 3, two-way ANOVA test, *P < 0.05, **P < 0.01, ***P < 0.001.

### Reduced VDR and enhanced bacteria in human colon cancer tissue

VDR expression was decreased in the AOM-DSS induced colon cancer model (Fig. 7a). We continued to explore VDR in human colorectal colon samples. Our data showed that increased Jak2 and STAT3 were associated with the reduction of intestinal VDR in human CRC intestines (Fig. 7b), suggesting that the JAK/STAT3 is upregulated in human CRC with protective VDR. Interestingly, we identified bacteria in human colorectal colon samples. FISH data showed that *Bacteroides fragilis* in tumors from patients with CRC (Fig. 7c).

**Fig. 7.**
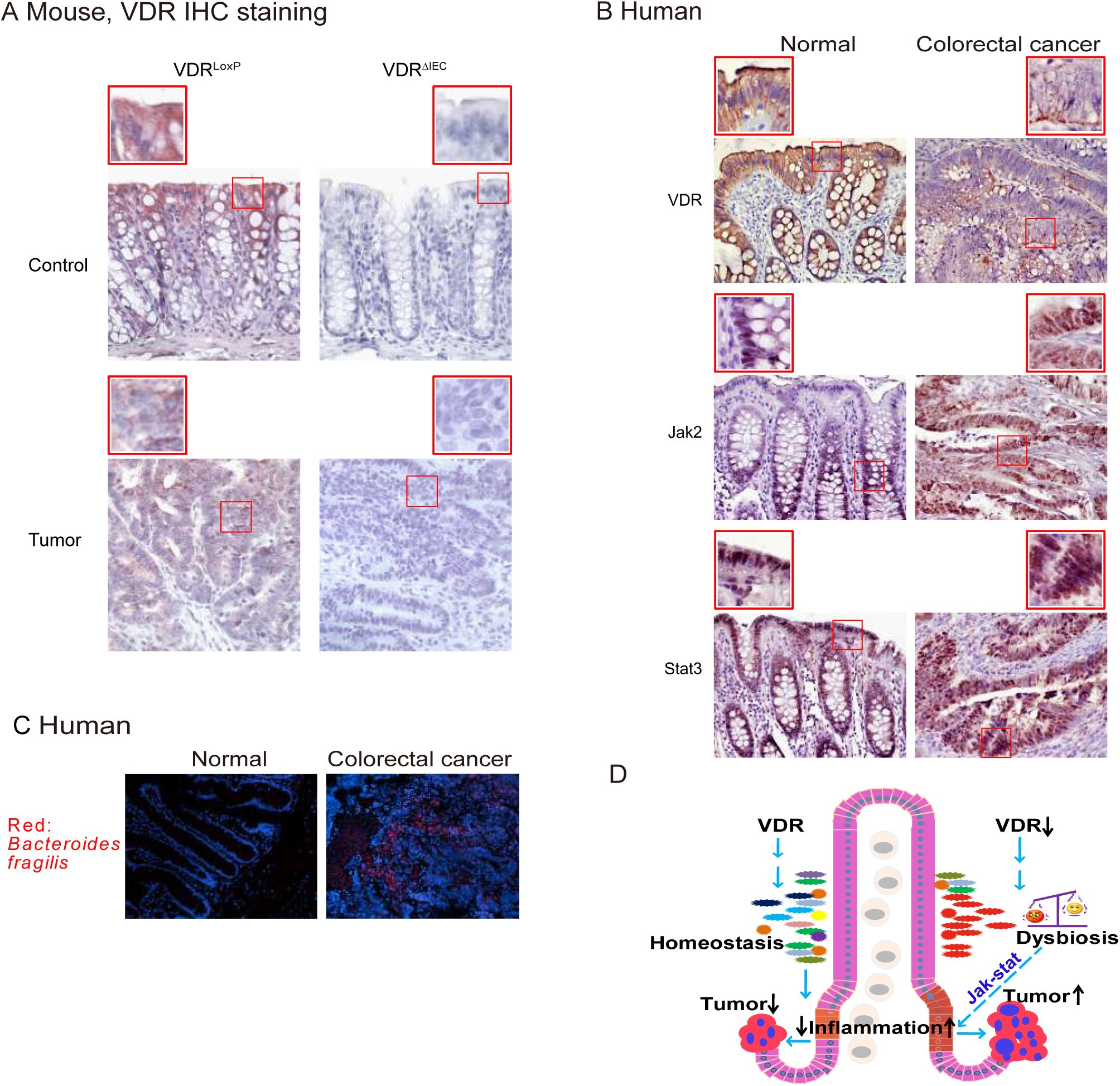
Enhanced bacteria, reduced VDR, increased Jak2 and STAT3 expression was observed in human CRC patients and AOM-DSS induced colon cancer model. **a** Intestinal VDR expression was decreased in the AOM-DSS induced colon cancer model. Images are from a single experiment and are representative of 6 mice per group. **b** Intestinal VDR, Jak2 and STAT3 staining in human CRC samples. Compared with normal intestines, CRC patients’ intestines had a statistically significantly lower VDR and higher Jak2/STAT3 expression. Images are representative of experiments that we carried out in triplicate; Normal, n=10; Colorectal cancer, n=10. **c** Bacteroides fragilis were found in human CRC samples compared to the normal tissue. Images are representative of experiments that we carried out in triplicate; Normal, n=10; Colorectal cancer, n=10. **d** A working model of intestinal VDR in regulating microbiome and colon cancer. Lack of VDR leads to dysbiosis and over-growth of tumors in colon.

## Discussion

In the current study, we have demonstrated that VDR deficiency in intestine leads to bacterial profile shift from normal to susceptible carcinogenesis. VDR^ΔIEC^ mice have higher tumor numbers with tumor location shifted from distal to proximal colon. Enhanced bacterial staining was found in tumors. Microbial metabolites from VDR^ΔIEC^ mice showed elevated secondary bile acids, which is consistent with the observations in human CRC. Furthermore, our study provides the mechanism of VDR dysfunction leading to dysbiosis and tumorigenesis through the hyperfunctioned Jak2. Fecal samples from VDR^ΔIEC^ mice enhance the STAT3 activation in human and mouse organoids. A JAK/STAT inhibitor abolished the microbiome-induced activation of STAT3. Our study fills the gaps by revealing mechanisms that are important to normal intestinal homeostasis and to chronic inflammation and dysbiosis, thus suggesting new therapeutic targets for restoring VDR functions in colitis-associated colon cancer (Fig. 7d working model).

Epidemiological and experimental studies have indicated a protective action of vitamin D against colorectal cancer [22–27]. Vitamin D_3_ exerts its chemopreventive activity by interrupting a crosstalk between tumor epithelial cells and the tumor microenvironment in a VDR-dependent manner [23]. Moreover, there is increasing interests regarding the use of vitamin D compounds for disease prevention and therapy [28]. If we do not understand the mechanism of the receptor of vitamin D, vitamin D taken by people may not be used effectively and efficiently. Hence, our current study fills the gap by characterizing the precise role for intestinal epithelial VDR in colon cancer models.

Endogenous enteric bacteria play a crucial role in the pathogenesis of colon cancer [29]. Dysregulation of bacterial-host interactions can result in chronic inflammatory and development of cancer [30, 31]. Multiple mechanisms of VDR affects cancers have been found, focusing on the host factors, e.g. beta-catenin pathway and inflammation [32]. However, very little is known about the physiological effects and molecular mechanisms responsible for intestinal epithelial VDR regulation of the microbiome community. Our study on VDR regulation of gut bacteria has demonstrated a microorganism-induced program of epithelial cell homeostasis and repair in the intestine [10, 33]. Abundance of Parabacteroides affected by VDR signaling in both human and mouse samples [8]. However, the specific relationship between the function of intestinal VDR and microbiome in tumorigenesis is not understood [34]. Here, we find out that VDR directly regulates host-bacterial interactions via JAK/STAT pathways and its downstream genes. Microbial metabolites from VDR^ΔIEC^ mice showed bile acid dysregulation and elevated secondary bile acids, which is consistent with the observed microbiome markers in human CRC [11, 12].

We used colonoids and mice lacking intestinal VDR expression to confirm the physiological relevance and molecular mechanism in epithelial-microbiome interactions. Research of intestinal VDR provides a framework to understand how the intestinal epithelial cells in the gut may inadvertently promote the development of cancer as an extension of its normal role in defense and repair. These insights are important for understanding health as well as disease.

We note a consistent link between low vitamin D/VDR signaling and high intestinal inflammation. Our studies suggest that cells lacking VDR are in a pre-inflammatory stat [10, 35, 36] and overexpression of VDR substantially reduced inflammation in VDR^−/−^ cells [35]. VDR is also identified as a suppressor of IFN-α-induced signaling through the JAK-STAT pathway [37]. The JAK/STAT pathway plays a critical role in intestinal and microbial homeostasis [38]. The JAK/STAT inhibitors have been recently tested as novel biological therapeutic strategies in inflammatory bowel diseases [15]. Because low dose proinflammatory cytokines are sufficient to induce bacterial endocytosis by epithelial cells, sub-clinical or low-grade changes below the threshold may tip the balance of tolerance towards full blown inflammation owing to subsequent intracellular microbial sensing and paracellular permeability damage. VDR expression increases epithelial integrity and attenuates inflammation. Thus, it is not surprising that the mucosal inflammation associated with VDR downregulation in intestine contributes to the initiation and progression of colon cancer.

## Conclusion

We provide a definitive characterization of the intestinal epithelial VDR in regulating diversity of the microbiome and colon cancer. It opens a new direction in the understanding of the microbial-VDR interactions in inflammation and caner. It indicates a new target — microbiome and VDR for prevention of cancer. VDR expression was decreased in the colon cancer mice after AOM/DSS treatment, which is consistent with the clinical observation in colitis-associated colon cancer patients [39]. In the future, we could also consider restoring the protective role of intestinal epithelia VDR using VDR activotrs or probiotics in CRC. Understanding of the abnormal interactions between host and microbiome will aid in developing novel strategies for managing chronic inflammatory diseases and cancers.

## Materials and Methods

### Human tissue samples

This study was performed in accordance with approval from the University of Rochester Ethics Committee (RSRB00037178). Colorectal tissue samples were obtained from 10 CRC patients with neoplasia and 10 patients without neoplasia patients (49–74years old). Human endoscopy samples in UIC hospital were collected for human organoids culture (IRB number 2017-0384).

### Animals

VDR^LoxP^ mice were originally reported by Dr. Geert Carmeliet [40]. VDR^ΔIEC^ mice were obtained by crossing the VDR^LoxP^ mice with villin-cre mice (Jackson Laboratory, 004586), as we previously reported [7]. Experiments were performed on 2–3 months old mice including male and female.

Mice were provided with water ad libitum and maintained in a 12 h dark/light cycle. The animal work was approved by the University of Rochester (When Dr. Sun’s lab was at University of Rochester), Rush University Animal Resources committee, and UIC Office of Animal Care.

### Induction of colon cancer by AOM-DSS in mice

Mice were treated with 10mg/kg of AOM (Sigma-Aldrich, Milwaukee, WI, USA) by intraperitoneal injection as previously described [41]. After a 7-day recovery period, mice received three cycles of 2% DSS in the drinking water. The initial sample size was 30 mice in the control group with no treatment and 30 in each experimental group. Tumor counts and measurements were performed in a blinded fashion under a stereo-dissecting microscope (Nikon SMZ1000, Melville, NY, USA). Microscopic analysis was performed for severity of inflammation and dysplasia on hematoxylin and eosin-stained ‘Swiss rolled’ colons by a gastrointestinal pathologist blinded to treatment conditions. Mice were scarified under anaesthesia.

### Cell culture

HCT116 cells were grown in high glucose Dulbecco’s Modified Eagle Medium (DMEM) (Hyclone, SH30243.01) containing 10% (v/v) fetal bovine serum (GEMINI, 900-108), 50 μg/ml streptomycin, and 50 U/ml penicillin (Mediatech, Inc., 30-002CI), as previously described [42, 43].

### Colonoids cultures and treatment with mice feces

C57BL/6J mice colonoids were prepared and maintained as previously described [44, 45]. Mini gut medium (advanced DMEM/F12 supplemented with HEPES, L-glutamine, N2, and B27) was added to the culture, along with R-Spondin, Noggin, EGF, and Wnt-3a. At day 7 after passage, colonoids were colonized by indicated mice feces for 2 hours, then washed, and incubated for 2 hours in Mini gut medium with Gentamicin (500 µg/ml).

Human organoids were developed using endoscopy samples in UIC hospital. Crypts were released from colon tissue by incubation for 30 min at 4 °C in PBS containing 2 mM EDTA. Isolated crypts were counted and pelleted. A total of 500 crypts were mixed with 50 μl of Matrigel (BD Bioscience) and plated in 24-well plates [46]. The colonoids were maintained in Human IntestiCult™ Organoid Growth Medium (STEMCELL Technologies Inc.).

Fresh feces were collected from 5 healthy VDR^LoxP^ or VDR^ΔIEC^ mice (8 weeks) and then well-mixed. 100mg feces homogenized in 6 ml Hanks and centrifuged for 30 s at 300 rpm, 4°C, to pellet the particulate matter. Organoids were treated with 250 μl feces supernatant for 2 hours, washed the organoids 3x with Hanks, and then incubated the cells in regular organoids culture medium for 2 hours [47].

### Western blot analysis and antibodies

Mouse colonic epithelial cells were collected by scraping the tissue from the colon of the mouse, including the proximal and distal regions [42, 48]. The cells were sonicated in lysis buffer (10 mM Tris, pH 7.4, 150 mM NaCl, 1 mM EDTA, 1 mM EGTA, pH 8.0, 1% Triton X-100) with 0.2 mM sodium ortho-vanadate, and protease inhibitor cocktail. The protein concentration was measured using the BioRad Reagent (BioRad, Hercules, CA, USA). Cultured cells were rinsed twice with ice-cold HBSS, lysed in protein loading buffer (50 mM Tris, pH 6.8, 100 mM dithiothreitol, 2% SDS, 0.1% bromophenol blue, 10% glycerol), and then sonicated. Equal amounts of protein were separated by SDS-polyacrylamide gel electrophoresis, transferred to nitrocellulose, and immunoblotted with primary antibodies. The following antibodies were used: anti-STAT3 (Cell Signaling Technology, 9132), anti-Jak2 (Cell Signaling Technology, 3230), anti-VDR (Santa Cruz Biotechnology, SC-13133), anti-Villin (Santa Cruz Biotechnology, SC-7672), anti-p-β-catenin (Cell Signaling Technology, 9566), anti-β-catenin (BD Biosciences, 610154), anti-PCNA (Santa Cruz Biotechnology, SC-25280), anti-LC3B (Cell Signaling Technology, 2775), or anti-β-actin (Sigma-Aldrich, A5316) antibodies and were visualized by ECL (Thermo Fisher Scientific, 32106). Membranes that were probed with more than one antibody were stripped before re-probing.

### Immunofluorescence

Colonic tissues were freshly isolated and embedded in paraffin wax after fixation with 10% neutral buffered formalin. Immunofluorescence was performed on paraffin-embedded sections (4 μm), after preparation of the slides as described previously [43] followed by incubation for 1 hour in blocking solution (2% bovine serum albumin, 1% goat serum in HBSS) to reduce nonspecific background. The tissue samples were incubated overnight with primary antibodies at 4°C. The following antibodies were used: anti-CD3, anti-CD11B and anti-CD68 (Santa Cruz Biotechnology), Slides were washed 3 times for 5 minutes each at room temperature in wash buffer. Samples were then incubated with secondary antibodies (goat anti-rabbit Alexa Fluor 488, Molecular Probes, CA; 1:200) for 1 hour at room temperature. Tissues were mounted with SlowFade Antifade Kit (Life technologies, s2828, Grand Island, NY, USA), followed by a coverslip, and the edges were sealed to prevent drying. Specimens were examined with a Zeiss laser scanning microscope LSM 710 (Carl Zeiss Inc., Oberkochen, Germany).

### Fluorescence in situ hybridization

Fluorescent in situ hybridization [49] was performed using antisense ssDNA probes targeting the bacterial 16S rRNA. Bfra602 probe (5’-GAGCCGCAAACTTTCACAA-3’) for Bacteroides fragilis group [50]. Prior to performing the FISH assay 5 μm tissue sections were baked over night at 55 °C. Tissue sections were deparaffinized in xylene, dehydrated with 100% ethanol, air dried, incubated in 0.2M HCl for 20min and heated in 1 mM sodium thiocyanate at 80 °C for 10 minutes. Samples were pepsin digested (4% pepsin in 0.01N HCl) for 20 minutes at 37 °C, washed slides in wash buffer (0.3 M NaCl, 0.03 M sodium citrate, pH 7, and 0. 1% SDS) and fixed the slides in 10% buffered formalin for 15min, washed and dried the slides, and hybridized with the probes at 5 ng/μl concentration each for 5 min at 96°C in hybridization buffer (0.9 M NaCl, 30% formamide, 20 mM Tris-HCl (pH 7.4), and 0.01% sodium dodecyl sulfate (SDS) and incubated at 37°C overnight. Slides were washed 4 times for 5 minutes each at 45°C in wash buffer. For visualization of the epithelial cell nuclei, the slides were counterstained with 4′, 6′-diamidino-2-phenylindole (DAPI) / antifade solution. Slides were examined with a Zeiss laser scanning microscope LSM 710 (Carl Zeiss Inc., Oberkochen, Germany).

### Mouse cytokines

Mouse blood samples were collected by cardiac puncture and placed in tubes containing EDTA (10 mg/ml). Mouse cytokines were measured using a mouse cytokine 10-Plex Panel kit (Invitrogen, Carlsbad, CA, USA) according to the manufacturer’s instructions. Briefly, beads of defined spectral properties were conjugated to protein-specific capture antibodies and added along with samples (including standards of known protein concentration, control samples, and test samples) into the wells of a filter-bottom microplate, where proteins bound to the capture antibodies over the course of a 2-hour incubation. After washing the beads, protein-specific biotinylated detector antibodies were added and incubated with the beads for 1 hour. After removal of excess biotinylated detector antibodies, the streptavidin-conjugated fluorescent protein R-phycoerythrin (streptavidin-RPE) was added and allowed to incubate for 30 minutes. After washing to remove unbound streptavidin-RPE, the beads were analyzed with the Luminex detection system (PerkinElmer CS1000 Autoplex Analyzer).

### Real Time quantitative PCR

Total RNA was extracted from epithelial cell monolayers or mouse colonic epithelial cells using TRIzol reagent (Thermo Fisher Scientific, 15596026). The RNA integrity was verified by gel electrophoresis. RNA reverse transcription was done using the iScript cDNA synthesis kit (Bio-Rad Laboratories, 1708891) according to the manufacturer’s directions. The RT-cDNA reaction products were subjected to quantitative real-time PCR using the MyiQ single-color real-time PCR detection system (Bio-Rad Laboratories, Hercules, CA, USA) and iTaq^TM^ Universal SYBR green supermix (Bio-Rad Laboratories, 1725121) according to the manufacturer’s directions. All expression levels were normalized to β-actin levels of the same sample. Percent expression was calculated as the ratio of the normalized value of each sample to that of the corresponding untreated control cells. All real-time PCR reactions were performed in triplicate. Primer sequences were designed using Primer-BLAST or were obtained from Primer Bank primer pairs listed in Table S1.

### Real-time PCR measurement of bacterial DNA

Mice feces samples DNA was extracted using stool DNA Kit (Omega bio-tek, Norcross, GA, USA) according to the manufacturer’s instructions. 16S rDNA PCR reactions were used the MyiQ single-color real-time PCR detection system (Bio-Rad Laboratories, Hercules, CA, USA) and iTaq^TM^ Universal SYBR green supermix (Bio-Rad Laboratories, 1725121) according to the manufacturer’s directions. Primers specific to 18S rRNAwere used as an endogenous control to normalize loading between samples [51]. The relative amount of 16S rDNA in each sample was estimated using the ΔΔCT. Primer sequences were designed using Primer-BLAST or were obtained from Primer Bank primer pairs listed in Table S2.

### Chromatin immunoprecipitation (CHIP) assay

Binding of VDR to the Jak2 promoter was investigated using the ChIP assay as described previously [36]. Briefly, HCT116 cells were treated with 1% formaldehyde for 10 min at 37°C. Cells were washed twice in ice-cold phosphate buffered saline containing protease inhibitor cocktail tablets (Roche). Cells were scraped into conical tubes, pelleted and lysed in SDS Lysis Buffer. The lysate was sonicated to shear DNA into fragments of 200–1000 bp (4 cycles of 10 s sonication, 10 s pausing, Branson Sonifier 250, USA). The chromatin samples were pre-cleared with salmon sperm DNA–bovine serum albumin-sepharose beads, then incubated overnight at 4 °C with VDR antibody (Santa Cruz Biotechnology). Immune complexes were precipitated with salmon sperm DNA–bovine serum albumin-sepharose beads. DNA was prepared by treatment with proteinase K, extraction with phenol and chloroform, and ethanol precipitation. Searching mouse ATG16L1 gene, we found a similar sequence as the VDRE sequence “(G/A)G(G/T)TCA”. We then designed primers for ChIP. PCR was performed using the following promoter specific primers: Jak2 forward, 5’-TGAATCCCAGGACACATTT-3’; reverse, 5’-GGTAAGCCACTGAAGGTT-3’.

### Histology of Intestine

Intestines were harvested, fixed in 10% formalin (pH 7.4), processed, and paraffin embedded. Sections (5μm) were stained with H&E. For immunostaining, antigens were retrieved by 10-minute boiling in 10 mM citrate (pH 6.0). The slides were stained with antibodies as previously described [43]. Blinded histological inflammatory scores were performed by a validated scoring system by a trained pathologist [52].

### LPS detection

LPS in serum samples was measured with Limulus amebocyte lysate (LAL) chromogenic endpoint assays (HIT302, Hycult Biotech, Plymouth Meeting, PA, USA) according to the manufacturer’s indications. The samples were diluted 1:4 with endotoxin-free water and then heated at 75°C for 5 min in a warm plate to denature the protein before the reaction. A standard curve was generated and used to calculate the concentrations, which were expressed as EU/ml, in the blood samples.

### Quantification of Fecal and Serum Lipocalin 2 (Lcn-2) by ELISA

Freshly collected fecal samples were reconstituted in PBS containing 0.1% Tween 20 (100 mg/ml) and vortexed for 20 min to get a homogenous fecal suspension. These samples were then centrifuged for 10 min at 12,000 rpm and 4°C. Clear supernatants were collected. Lcn-2 levels were estimated in the supernatants using Duoset murine Lcn-2 ELISA kit (R&D Systems, Minneapolis, MN), as described in our previous study [53].

### Mucosa microbial and fecal 454 Pyrosequencing

The tubes for microbial sampling were autoclaved and then irradiated with ultraviolet light to destroy the environmental bacterial DNA. The mice were then anesthetized and dissected. Fecal isolated freshly from the gut and placed into the specially prepared tubes, as described in our previously published papers [54, 55]. The samples were kept at low temperature with dry ice and mailed to Research and Testing Laboratory, Lubbock, TX, for 454 pyrosequencing. The V4-V6 region of the samples was amplified in Research and Testing Laboratory, Lubbock, TX, for pyrosequencing using a forward and reverse fusion primer. The sequences were denoised, subjected to quality checking. Taxonomic identifications were assigned by queries against NCBI.

### Sample Preparation for Metabolites

Fecal samples were prepared using the automated MicroLab STAR® system from Hamilton Company. Several recovery standards were added prior to the first step in the extraction process for QC purposes. To remove protein, dissociate small molecules bound to protein or trapped in the precipitated protein matrix, and to recover chemically diverse metabolites, proteins were precipitated with methanol under vigorous shaking for 2 min (Glen Mills GenoGrinder 2000) followed by centrifugation. The resulting extract was divided into five fractions: two for analysis by two separate reverse phase (RP)/UPLC-MS/MS methods with positive ion mode electrospray ionization (ESI), one for analysis by RP/UPLC-MS/MS with negative ion mode ESI, one for analysis by HILIC/UPLC-MS/MS with negative ion mode ESI, and one sample was reserved for backup. Samples were placed briefly on a TurboVap® (Zymark) to remove the organic solvent. The sample extracts were stored overnight under nitrogen before preparation for analysis.

### Metabolite analysis

For metabolite experiment, 33 of divided into VDR^ΔIEC^ (N= 17) and control VDR^LoxP^ (N=16) groups. All mice were housed in specific pathogen-free environments under a controlled condition of 12 h light/12 h dark cycle at 20–22 °C and 45 ± 5% humidity, with free access to food and ultrapure water. At 16 weeks of age fecal contents of each mouse were carefully collected in separate Eppendorf tubes, labeled with a unique identification number and stored at −80 °C until shipped. Samples were transported to Metabolon Inc, NC, USA in dry ice by overnight shipment for analysis.

Following receipt, samples were assigned a unique identifier by the LIMS (laboratory information management system) and immediately stored at −80°C until processed. Samples were prepared using the automated MicroLab STAR® system from Hamilton Company. First proteins and other associated small molecules were precipitated then diverse metabolites were recovered by grinding and centrifugation. The resulting extract was analyzed by two separate reverse phase (RP)/UPLC-MS/MS methods with positive ion mode electrospray ionization (ESI), or with negative ion mode ESI, and one by HILIC/UPLC-MS/MS with negative ion mode ESI. Several types of controls were analyzed along with the experimental samples to ensure accurate and consistent identification. Ultrahigh Performance Liquid Chromatography-Tandem Mass Spectroscopy (UPLC-MS/MS) was utilized as an analyzer. Metabolon’s hardware and software were used to extract the raw data, followed by the identification of peak and QC (Quality Check). These systems are built on a web-service platform utilizing Microsoft’s NET technologies.

### Microbiome data analysis

Differences in microbial communities between VDR^LoxP^ and VDR^ΔIEC^ groups were analyzed, as we did in previous studies [54, 55]. Briefly, Principal Coordinates Analysis (PCoA) of unweighted UniFrac distances plots were plotted using quantitative insights into microbial ecology (QIIME) [56]. To determine differences in microbiota composition between the animal groups, the analysis of similarities (ANOSIM) function in the statistical software package PRIMER 6 (PRIMER-E Ltd., Lutton, UK) was used on the unweighted UniFrac distance matrixes [57].

### Statistical Analysis

Metabolite data were expressed as fold change ratio, all other data are expressed as the mean ± SD. All statistical tests were 2-sided. The p values <0.05 were considered statistically significant. For metabolite data, following log transformation and imputation of missing values, if any, with the minimum observed value for each compound, ANOVA contrasts and Welch’s two-sample *t*-test were used to identify biochemicals that differed significantly between experimental groups. For other data analyses, the differences between samples were analyzed using Student’s *t*-test for two-groups comparison, one-way ANOVA for more than two-groups comparison, respectively.

## Declarations

### Ethics approval and consent to participate

The animal work was approved by the University of Rochester (When Dr. Sun’s lab was at University of Rochester), Rush University Animal Resources committee, and UIC Office of Animal Care.

Consent to participate is not applicable.

### Consent for publication

Not applicable.

### Availability of data and material

Data and materials are available upon request.

### Authors’ contributions

YZ, RL, SW: acquisition, analysis and interpretation of data; drafting of the manuscript; statistical analysis. IC: metabolite data analysis. DZ: Pathological, technical and material support. YX: statistical analysis, microbiome data analysis, and drafting of the manuscript. JS: study concept and design; analysis and interpretation of data; writing the manuscript for important intellectual content, obtained funding, and study supervision.

## Acknowledgements

We would like to thank Eric Xia and Jason Xia for helping with proofreading.

## Availability of supporting data

Sequence files and metadata for all samples used in this study have been deposited in https://www.ncbi.nlm.nih.gov/bioproject/593562

SubmissionID: SUB6615727

BioProject ID: PRJNA593562

## Abbreviations

1,25(OH)_2_D_3_: 1α,25-dihydroxy vitamin D_3_
AOM: azoxymethane
BrdU: bromodeoxyuridine
CHIP: Chromatin immunoprecipitation
CRC: colon rectal cancer
DSS: dextran sodium sulfate
FISH: Fluorescent in situ hybridization
IECs: Intestinal epithelial cells
Lcn-2: Lipocalin 2
IL10: Interleukin 10
Jak: Janus kinases
LPS: Lipopolysaccharides
PCNA: Proliferating cell nuclear antigen
STAT3: Signal transducer and activator of transcription 3
TUNEL: terminal transferase-mediated dUTP nick end labeling
VDR: vitamin D receptor

---

**Sup. Table 1.**
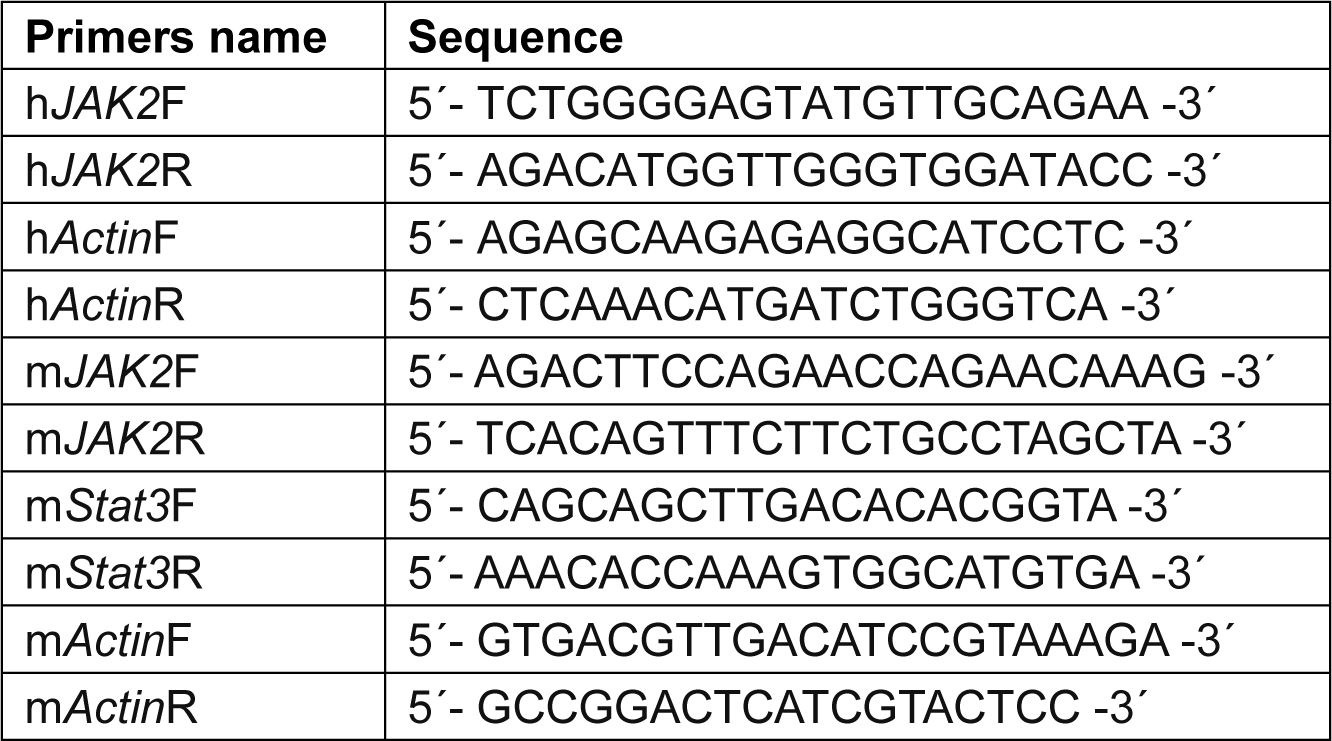
Real-time PCR primers.

**Sup. Table 2.**
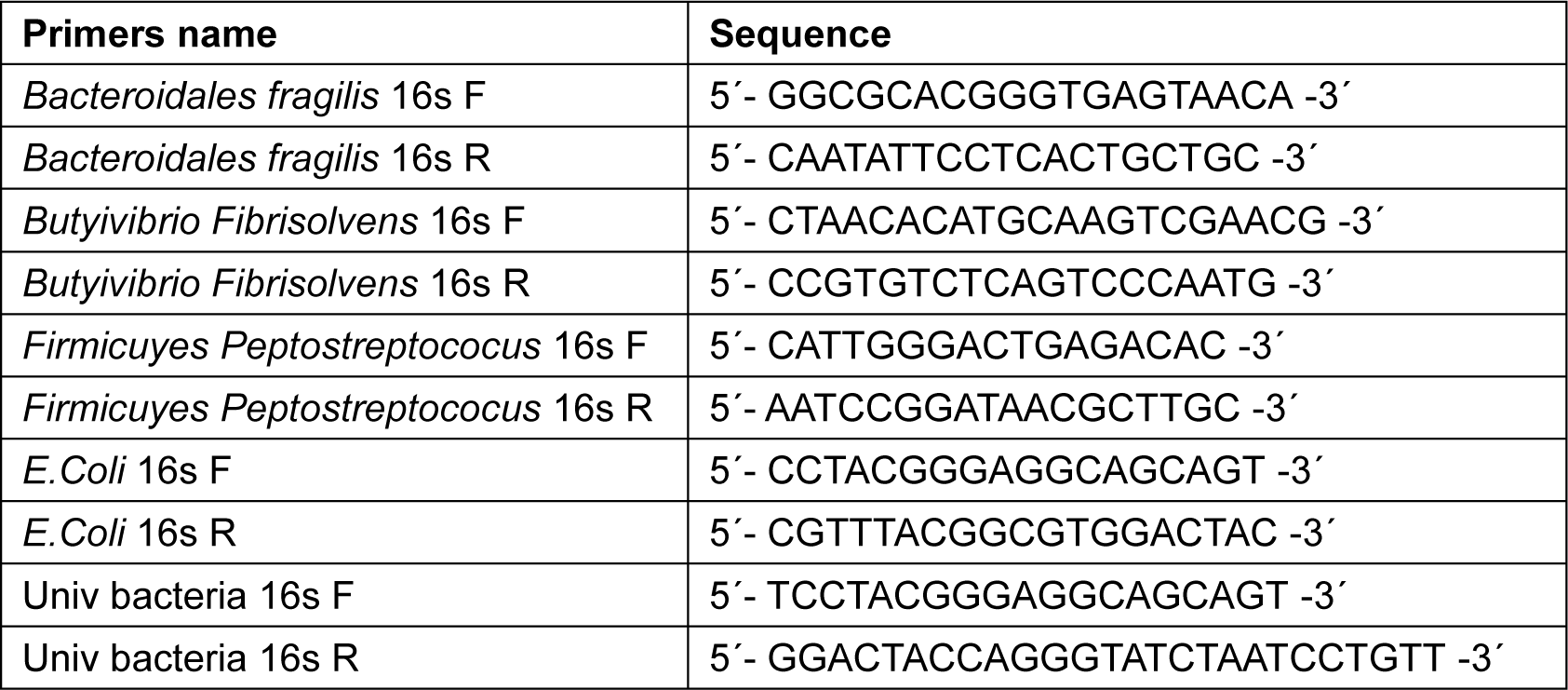
Bacterial 16S rDNA Real-time PCR primers.

## References

1. McCullough ML, Zoltick ES, Weinstein SJ, Fedirko V, Wang M, Cook NR, Eliassen AH, Zeleniuch-Jacquotte A, Agnoli C, Albanes D, et al: Circulating Vitamin D and Colorectal Cancer Risk: An International Pooling Project of 17 Cohorts. J Natl Cancer Inst 2018.

2. Ferrer-Mayorga G, Gomez-Lopez G, Barbachano A, Fernandez-Barral A, Pena C, Pisano DG, Cantero R, Rojo F, Munoz A, Larriba MJ: Vitamin D receptor expression and associated gene signature in tumour stromal fibroblasts predict clinical outcome in colorectal cancer. Gut 2017, 66:1449–1462.

3. Vaughan-Shaw PG, Zgaga L, Ooi LY, Theodoratou E, Timofeeva M, Svinti V, Walker M, O’Sullivan F, Ewing A, Johnston S, et al: Low plasma vitamin D is associated with adverse colorectal cancer survival after surgical resection, independent of systemic inflammatory response. Gut 2019:1–9.

4. Ng K, Nimeiri HS, McCleary NJ, Abrams TA, Yurgelun MB, Cleary JM, Rubinson DA, Schrag D, Miksad R, Bullock AJ, et al: Effect of High-Dose vs Standard-Dose Vitamin D3 Supplementation on Progression-Free Survival Among Patients With Advanced or Metastatic Colorectal Cancer: The SUNSHINE Randomized Clinical Trial. JAMA 2019, 321:1370–1379.

5. Haussler MR, Whitfield GK, Haussler CA, Hsieh J-C, Thompson PD, Selznick SH, Dominguez CE, Jurutka PW: The nuclear vitamin D receptor: biological and molecular regulatory properties revealed. J Bone and Mineral Research 1998, 13:325–349.

6. Thorne JaC, J. M.: The molecular cell biology of VDR. Springer Science & Business Media; 2010.

7. Wu S, Zhang YG, Lu R, Xia Y, Zhou D, Petrof EO, Claud EC, Chen D, Chang EB, Carmeliet G, Sun J: Intestinal epithelial vitamin D receptor deletion leads to defective autophagy in colitis. Gut 2015, 64:1082–1094.

8. Wang J, Thingholm LB, Skieceviciene J, Rausch P, Kummen M, Hov JR, Degenhardt F, Heinsen FA, Ruhlemann MC, Szymczak S, et al: Genome-wide association analysis identifies variation in vitamin D receptor and other host factors influencing the gut microbiota. Nat Genet 2016, 48:1396–1406.

9. Sun J: VDR/vitamin D receptor regulates autophagic activity through ATG16L1. Autophagy 2016, 12:1057–1058.

10. Wu S, Liao AP, Xia Y, Li YC, Li JD, Sartor RB, Sun J: Vitamin D receptor negatively regulates bacterial-stimulated NF-kappaB activity in intestine. Am J Pathol 2010, 177:686–697.

11. Thomas AM, Manghi P, Asnicar F, Pasolli E, Armanini F, Zolfo M, Beghini F, Manara S, Karcher N, Pozzi C, et al: Metagenomic analysis of colorectal cancer datasets identifies cross-cohort microbial diagnostic signatures and a link with choline degradation. Nat Med 2019, 25:667–678.

12. Wirbel J, Pyl PT, Kartal E, Zych K, Kashani A, Milanese A, Fleck JS, Voigt AY, Palleja A, Ponnudurai R, et al: Meta-analysis of fecal metagenomes reveals global microbial signatures that are specific for colorectal cancer. Nat Med 2019, 25:679–689.

13. Buchert M, Burns CJ, Ernst M: Targeting JAK kinase in solid tumors: emerging opportunities and challenges. Oncogene 2016, 35:939–951.

14. Lu R, Zhang YG, Sun J: STAT3 activation in infection and infection-associated cancer. Mol Cell Endocrinol 2017, 451:80–87.

15. D’Amico F, Fiorino G, Furfaro F, Allocca M, Danese S: Janus kinase inhibitors for the treatment of inflammatory bowel diseases: developments from phase I and phase II clinical trials. Expert Opin Investig Drugs 2018, 27:595–599.

16. Bird RP, Good CK: The significance of aberrant crypt foci in understanding the pathogenesis of colon cancer. Toxicol Lett 2000, 112-113:395–402.

17. Gerard P: Metabolism of cholesterol and bile acids by the gut microbiota. Pathogens 2013, 3:14–24.

18. Sagar NM, Cree IA, Covington JA, Arasaradnam RP: The interplay of the gut microbiome, bile acids, and volatile organic compounds. Gastroenterol Res Pract 2015, 2015:398585.

19. Staley C, Weingarden AR, Khoruts A, Sadowsky MJ: Interaction of gut microbiota with bile acid metabolism and its influence on disease states. Appl Microbiol Biotechnol 2017, 101:47–64.

20. Li T, Apte U: Bile Acid Metabolism and Signaling in Cholestasis, Inflammation, and Cancer. Adv Pharmacol 2015, 74:263–302.

21. Chassaing B, Srinivasan G, Delgado MA, Young AN, Gewirtz AT, Vijay-Kumar M: Fecal lipocalin 2, a sensitive and broadly dynamic non-invasive biomarker for intestinal inflammation. PLoS One 2012, 7:e44328.

22. Protiva P, Cross HS, Hopkins ME, Kallay E, Bises G, Dreyhaupt E, Augenlicht L, Lipkin M, Lesser M, Livote E, Holt PR: Chemoprevention of colorectal neoplasia by estrogen: potential role of vitamin D activity. Cancer Prev Res (Phila Pa) 2009, 2:43–51.

23. Kaler P, Augenlicht L, Klampfer L: Macrophage-derived IL-1beta stimulates Wnt signaling and growth of colon cancer cells: a crosstalk interrupted by vitamin D3. Oncogene 2009, 28:3892–3902.

24. Fichera A, Little N, Dougherty U, Mustafi R, Cerda S, Li YC, Delgado J, Arora A, Campbell LK, Joseph L, et al: A vitamin D analogue inhibits colonic carcinogenesis in the AOM/DSS model. J Surg Res 2007, 142:239–245.

25. Nagpal S, Lu J, Boehm MF: Vitamin D analogs: mechanism of action and therapeutic applications. Curr Med Chem 2001, 8:1661–1679.

26. Palmer HG, Sanchez-Carbayo M, Ordonez-Moran P, Larriba MJ, Cordon-Cardo C, Munoz A: Genetic signatures of differentiation induced by 1alpha,25-dihydroxyvitamin D3 in human colon cancer cells. Cancer Res 2003, 63:7799–7806.

27. Chan AT, Giovannucci EL: Primary prevention of colorectal cancer. Gastroenterology 2010, 138:2029–2043 e2010.

28. Gombart AF, Luong QT, Koeffler HP: Vitamin D compounds: activity against microbes and cancer. Anticancer Res 2006, 26:2531–2542.

29. Wong SH, Zhao L, Zhang X, Nakatsu G, Han J, Xu W, Xiao X, Kwong TNY, Tsoi H, Wu WKK, et al: Gavage of Fecal Samples From Patients With Colorectal Cancer Promotes Intestinal Carcinogenesis in Germ-Free and Conventional Mice. Gastroenterology 2017, 153:1621–1633 e1626.

30. Terzic J, Grivennikov S, Karin E, Karin M: Inflammation and colon cancer. Gastroenterology, 138:2101–2114 e2105.

31. Song M, Chan AT, Jun J: Features of the Gut Microbiome, Diet, and Environment That Influence Risk of Colorectal Cancer. Gastroenterology 2020.

32. Sun J: The Role of Vitamin D and Vitamin D Receptors in Colon Cancer. Clin Transl Gastroenterol 2017, 8:e103.

33. Abreu MT: Toll-like receptor signalling in the intestinal epithelium: how bacterial recognition shapes intestinal function. Nat Rev Immunol 2010, 10:131–144.

34. De Mattia E, Cecchin E, Montico M, Labriet A, Guillemette C, Dreussi E, Roncato R, Bignucolo A, Buonadonna A, D’Andrea M, et al: Association of STAT-3 rs1053004 and VDR rs11574077 With FOLFIRI-Related Gastrointestinal Toxicity in Metastatic Colorectal Cancer Patients. Front Pharmacol 2018, 9:367.

35. Sun J, Kong J, Duan Y, Szeto FL, Liao A, Madara JL, Li YC: Increased NF-kappaB activity in fibroblasts lacking the vitamin D receptor. Am J Physiol Endocrinol Metab 2006, 291:E315–322.

36. Wu S, Xia Y, Liu X, Sun J: Vitamin D receptor deletion leads to reduced level of IkappaBalpha protein through protein translation, protein-protein interaction, and post-translational modification. Int J Biochem Cell Biol 2010, 42:329–336.

37. Lange CM, Gouttenoire J, Duong FH, Morikawa K, Heim MH, Moradpour D: Vitamin D receptor and Jak-STAT signaling crosstalk results in calcitriol-mediated increase of hepatocellular response to IFN-alpha. J Immunol 2014, 192:6037–6044.

38. Heneghan AF, Pierre JF, Kudsk KA: JAK-STAT and intestinal mucosal immunology. JAKSTAT 2013, 2:e25530.

39. Wada K, Tanaka H, Maeda K, Inoue T, Noda E, Amano R, Kubo N, Muguruma K, Yamada N, Yashiro M, et al: Vitamin D receptor expression is associated with colon cancer in ulcerative colitis. Oncol Rep 2009, 22:1021–1025.

40. Van Cromphaut SJ, Dewerchin M, Hoenderop JG, Stockmans I, Van Herck E, Kato S, Bindels RJ, Collen D, Carmeliet P, Bouillon R, Carmeliet G: Duodenal calcium absorption in vitamin D receptor-knockout mice: functional and molecular aspects. Proc Natl Acad Sci U S A 2001, 98:13324–13329.

41. Greten FR, Karin M: The IKK/NF-kappaB activation pathway-a target for prevention and treatment of cancer. Cancer Lett 2004, 206:193–199.

42. Sun J, Hobert ME, Duan Y, Rao AS, He TC, Chang EB, Madara JL: Crosstalk between NF-kappaB and beta-catenin pathways in bacterial-colonized intestinal epithelial cells. Am J Physiol Gastrointest Liver Physiol 2005, 289:129–137.

43. Lu R, Wu S, Liu X, Xia Y, Zhang YG, Sun J: Chronic effects of a Salmonella type III secretion effector protein AvrA in vivo. PLoS One 2010, 5:e10505.

44. Zhang YG, Wu S, Xia Y, Sun J: Salmonella-infected crypt-derived intestinal organoid culture system for host-bacterial interactions. Physiol Rep 2014, 2.

45. Lu R, Voigt RM, Zhang Y, Kato I, Xia Y, Forsyth CB, Keshavarzian A, Sun J: Alcohol Injury Damages Intestinal Stem Cells. Alcohol Clin Exp Res 2017, 41:727–734.

46. Sato T, Vries RG, Snippert HJ, van de Wetering M, Barker N, Stange DE, van Es JH, Abo A, Kujala P, Peters PJ, Clevers H: Single Lgr5 stem cells build crypt-villus structures in vitro without a mesenchymal niche. Nature 2009, 459:262–265.

47. Zhang YG, Zhu X, Lu R, Messer JS, Xia Y, Chang EB, Sun J: Intestinal epithelial HMGB1 inhibits bacterial infection via STAT3 regulation of autophagy. Autophagy 2019.

48. Duan Y, Liao AP, Kuppireddi S, Ye Z, Ciancio MJ, Sun J: beta-Catenin activity negatively regulates bacteria-induced inflammation. Lab Invest 2007, 87:613–624.

49. Barrett JC, Hansoul S, Nicolae DL, Cho JH, Duerr RH, Rioux JD, Brant SR, Silverberg MS, Taylor KD, Barmada MM, et al: Genome-wide association defines more than 30 distinct susceptibility loci for Crohn’s disease. Nat Genet 2008, 40:955–962.

50. Franks AH, Harmsen HJ, Raangs GC, Jansen GJ, Schut F, Welling GW: Variations of bacterial populations in human feces measured by fluorescent in situ hybridization with group-specific 16S rRNA-targeted oligonucleotide probes. Appl Environ Microbiol 1998, 64:3336–3345.

51. Fan Y, Dickman KG, Zong WX: Akt and c-Myc differentially activate cellular metabolic programs and prime cells to bioenergetic inhibition. J Biol Chem 2010, 285:7324–7333.

52. Sellon RK, Tonkonogy S, Schultz M, Dieleman LA, Grenther W, Balish E, Rennick DM, Sartor RB: Resident enteric bacteria are necessary for development of spontaneous colitis and immune system activation in interleukin-10-deficient mice. Infect Immun 1998, 66:5224–5231.

53. Lu R, Zhang YG, Xia Y, Sun J: Imbalance of autophagy and apoptosis in intestinal epithelium lacking the vitamin D receptor. FASEB J 2019:fj201900727R.

54. Wang Y, Hoenig JD, Malin KJ, Qamar S, Petrof EO, Sun J, Antonopoulos DA, Chang EB, Claud EC: 16S rRNA gene-based analysis of fecal microbiota from preterm infants with and without necrotizing enterocolitis. ISME J 2009, 3:944–954.

55. Devkota S, Wang Y, Musch MW, Leone V, Fehlner-Peach H, Nadimpalli A, Antonopoulos DA, Jabri B, Chang EB: Dietary-fat-induced taurocholic acid promotes pathobiont expansion and colitis in Il10−/− mice. Nature 2012, 487:104–108.

56. Lozupone C, Knight R: UniFrac: a new phylogenetic method for comparing microbial communities. Appl Environ Microbiol 2005, 71:8228–8235.

57. Xia Y, Sun J, Chen D-G: Statistical analysis of microbiome data with R. Singapore: Springer 2018.

